# Post-Hurricane Vital Statistics Expose Fragility of Puerto Rico’s Health System

**DOI:** 10.1101/407874

**Authors:** Rolando J. Acosta, Rafael A. Irizarry

## Abstract

**Importance:** Hurricane Maria made landfall in Puerto Rico on September 20, 2017. As recently as May of this year (2018), the official death count was 64. After a study describing a household survey reported a much higher death count estimate, as well as evidence of population displacement, extensive loss of services, and a prolonged death rate the government released death registry data. These newly released data will permit a better understanding of the effects of this hurricane.

**Objective:** Provide a detailed description of the effects on mortality of Hurricane Maria and compare to other hurricanes.

**Design:** We fit a statistical model to mortality data that accounts for seasonal and non-hurricane related yearly effects. We then estimated the deviation from the expected death rate as a function of time.

**Setting:** We fit this model to 1985-2018 Puerto Rico daily data, which includes the dates of hurricanes Hugo, Georges, and Maria, 2015-2018 Florida daily data, which includes the dates of Hurricane Irma, 2002-2004 Louisiana monthly data, which includes the date of Hurricane Katrina, and 2000-2016 New Jersey monthly data, which includes the date of Hurricane Sandy.

**Results:** We find a prolonged increase in death rate after Maria and Katrina, lasting at least 207 and 125 days, resulting in excess deaths estimates of 3,400 (95% CI, 3,100-3,700), and 1,800 (95% CI, 1,600-2100) respectively, showing that Maria had a more long term damaging impact. Surprisingly, we also find that in 1998, Georges had a comparable impact to Katrina’s with a prolonged increase of 106 days resulting in 1,400 (95% CI, 1,200-1,700) excess deaths. For Hurricane Maria, we find sharp increases in a small number of causes of deaths, including diseases of the circulatory, endocrine and respiratory system, as well as bacterial infections and suicides.

**Conclusion and Relevance:** Our analysis suggests that since at least 1998, Puerto Rico’s health system has been in a precarious state. Without a substantial intervention, it appears that if hit with another strong hurricane, Puerto Ricans will suffer the unnecessary death of hundreds of its citizens.

**Key Points:** **Question:** How does the effect of Hurricane Maria on mortality in Puerto Rico compare to the effect of other hurricanes in Puerto Rico and other United States jurisdictions?

**Findings:** We estimate about 3,000 excess deaths after Maria, a higher toll than Katrina. Only other comparable effect was after Georges, also in Puerto Rico. For Georges and Maria, we observe a prolonged death rate increase of more than 10% lasting several months. The causes of death that increased after Maria are consistent with a collapsed health system

**Meaning:** Puerto Rico’s health system does not appear to be ready to withstand another strong hurricane.

## Introduction

Hurricane Maria made landfall in Puerto Rico on September 20, 2017, interrupting the water supply, electricity, telecommunications networks, and access to medical care. In early May 2018, the official death count stood at 64^1^. This figure was in conflict with estimates obtained by groups that ostensibly had access to death counts for September and October from the demographic registry of Puerto Rico. By comparing these two numbers to historical averages, additional deaths were estimated to be in excess of 1,000^2-5^. However, as of May 2017, the government of Puerto Rico was not releasing the 2017-2018 data.

On May 29, 2018 a multi-institutional study was published^6^, here referred to as the *Harvard Study,* describing a survey of 3,299 households and reporting a death count estimate of 4,645 (95% CI, 793-8,498). It also reported an extensive loss of services after the hurricane. Perhaps most importantly, the study showed evidence of a sustained increased effect on mortality throughout this extended period. These findings underscored the importance of a careful analysis to determine if, for example, there was a systematic increase in deaths due to indirect effects, if a specific demographic was at greater risk, and what type of medical conditions needed most attention.

The Harvard study received worldwide media coverage and three days after its publication, while under significant public pressure and facing a lawsuit^7^, the government finally made the data public and acknowledged the possibility of a higher death count of 1,427^8^. This number is consistent with the value one obtains by subtracting the 2017 monthly counts from the 2013-2016 average using the newly released data (Supplementary Table 1): (2928-2399)+(3040- 2514)+(2671-2418)+(2820-2701)=1427. Santos-Lozada and Howard^9^ updated their previous estimate to 1,139 (95% CI, 1,006-1,272) for September, October and November. However, neither of these estimates took into account the population displacement described by the Harvard study and others^10,11^. An analysis, posted online, took into account population displacement and suggested a larger count^12^. A government commissioned study came to a similar conclusion and provided an estimate of 2,975 (95% CI:2,658-3,290) for the total study period of September 2017 through February 2018^13^.

**Table 1:** Excess deaths by cause of death for categories with p-values below 0.05/30

| ICD | Observed | Expected | Excess | 95% CI |
| --- | --- | --- | --- | --- |
| [I00,I99] | 2511 | 1808 | 703 | [2,389-2,639] |
| [E00,E35] | 1118 | 788 | 330 | [1,037-1,205] |
| [G00,G99] | 961 | 672 | 289 | [886-1,042] |
| [J00,J99] | 898 | 629 | 269 | [826-976] |
| [A00,A79] | 362 | 201 | 161 | [315-416] |
| [V01,Y98] | 632 | 498 | 134 | [573-697] |
| [R95,R99] | 268 | 147 | 121 | [228-315] |
| [N00,N39] | 435 | 331 | 104 | [386-490] |
| [E70,E90] | 174 | 106 | 68 | [143-211] |
| [L00,L99] | 102 | 53 | 49 | [78-133] |
| [F00,F99] | 167 | 119 | 48 | [138-203] |
| Storm* | 34 | 0 | 34 | [5-248] |
| Suicide* | 81 | 48 | 33 | [61-108] |
| [M00,M99] | 78 | 46 | 32 | [58-105] |
| Leptospirosis* | 21 | 4 | 17 | [10-45] |

\* We combined all the causes of death descriptions with these words

Here we use these daily counts and individual level mortality data to provide a detailed picture of the effect Hurricane Maria had on mortality in Puerto Rico. We compare the death rate increases to those observed in Hugo and Georges, two previous hurricanes in Puerto Rico, Katrina in Louisiana, Irma in Florida, and Sandy in New Jersey. We find a disturbing pattern in the Puerto Rico data.

## Methods

### Data

We set out to obtain detailed mortality and population size data related to hurricanes Hugo, Georges, and Maria in Puerto Rico, Katrina in Louisiana, Sandy in New Jersey, Harvey in Texas, and Irma in Florida. We requested individual death information, but this was not always available. We used whatever data was made available, which resulted in a mix of individual, daily and monthly data.

### Hurricanes Hugo, Georges, and Maria (Puerto Rico)

We requested daily death count data from the Department of Health of Puerto Rico and obtained data from January 1985 to December 2014. We also requested individual level information with no personal identifiers from the Department of Health of Puerto Rico and obtained individual records including date, gender, age, and causes of death from January 2015 to June 2018. We used these data to construct the daily counts for the 2015-2018 period. Exploratory data analysis showed that data were incomplete after May 31, 2018 (Supplementary Figure 1) and discarded data past this date. Yearly population estimates for the island were obtained from the Puerto Rico Statistical Institute. We computed daily population estimates via linear interpolation (Supplementary Figure 2A). To obtain an estimate of the population displacement after Hurricane Maria, we used population movement estimates from Teralytics Inc.^14^ (Supplementary Figure 2B). We combined these two datasets to obtain a final estimate of the population (Supplementary Figure 2C).

**Figure 1:**
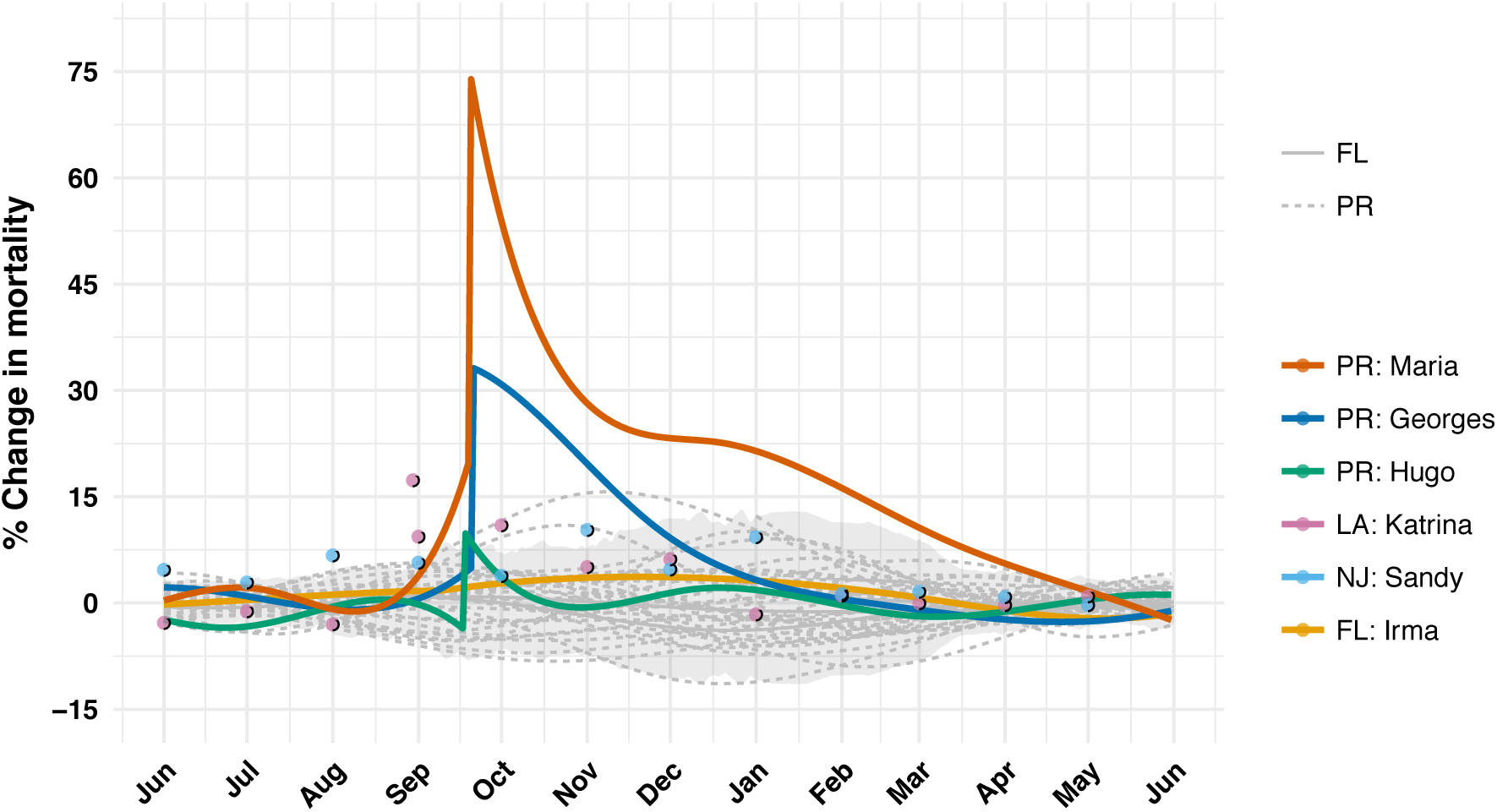
Estimated percent increases in death rates after hurricanes Hugo, Georges, and María in Puerto Rico, Katrina in Louisiana, Sandy in New Jersey and Irma in Florida. Estimates for Puerto Rico and Florida were obtained

**Figure 2:**
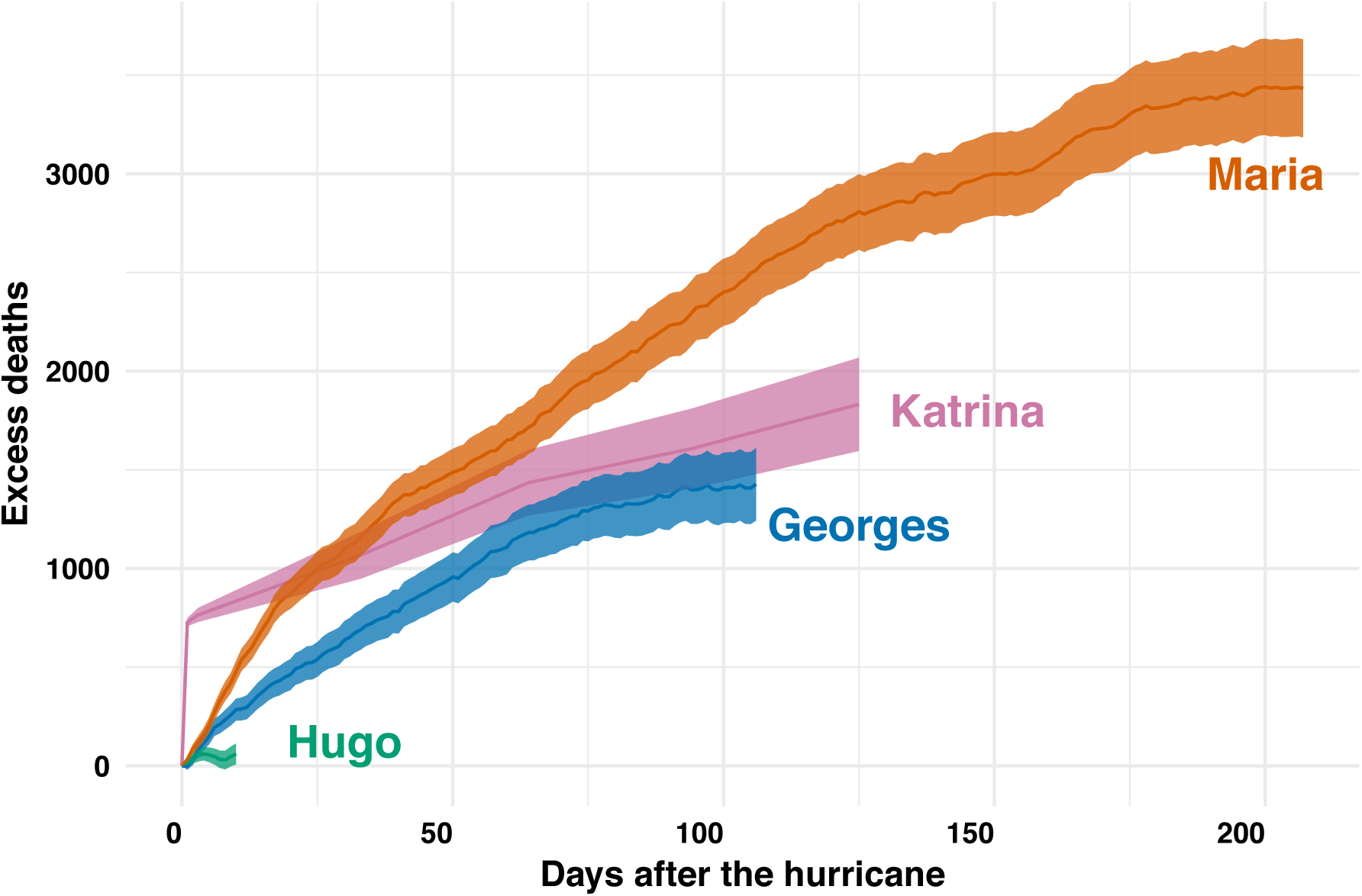
Excess deaths versus days after the hurricane for hurricanes Katrina, Georges, María, Hugo and Katrina. The shaded areas represent marginal 95% confidence intervals for these estimates. The excess deaths are calculated for the periods affected by indirect effects

### Hurricane Irma (Florida)

We requested daily death counts from Florida’s Vital Statistic System and obtained data from 2015 to 2018. For consistency, we discarded data past May 31, 2018. Yearly population estimates were obtained from the US Census for 2015-2017. We computed daily population estimates using interpolation and extrapolated using a linear model for 2018. Furthermore, we used data provided by Teralytics Inc. to estimate changes in population due to immigration from Puerto Ricans due to Hurricane Maria (Supplementary Figure 2D).

### Hurricane Katrina (Louisiana)

We requested daily death counts from Louisiana’s Vital Statistic System and obtained data from 2003-2006. For Louisiana, we also obtained monthly death counts from the Underlying Cause of Death database through CDC WONDER for 2000-2008^15^. These two datasets did not match for the months following Hurricane Katrina (Supplementary Figure 3). Since the data for August 2005 matched, we used the daily data to divide the monthly counts for August into before, during, and after the hurricane counts. We obtained population estimates from the US Census and computed daily population estimates via linear interpolation (Supplementary Figure 2E).

**Figure 3:**
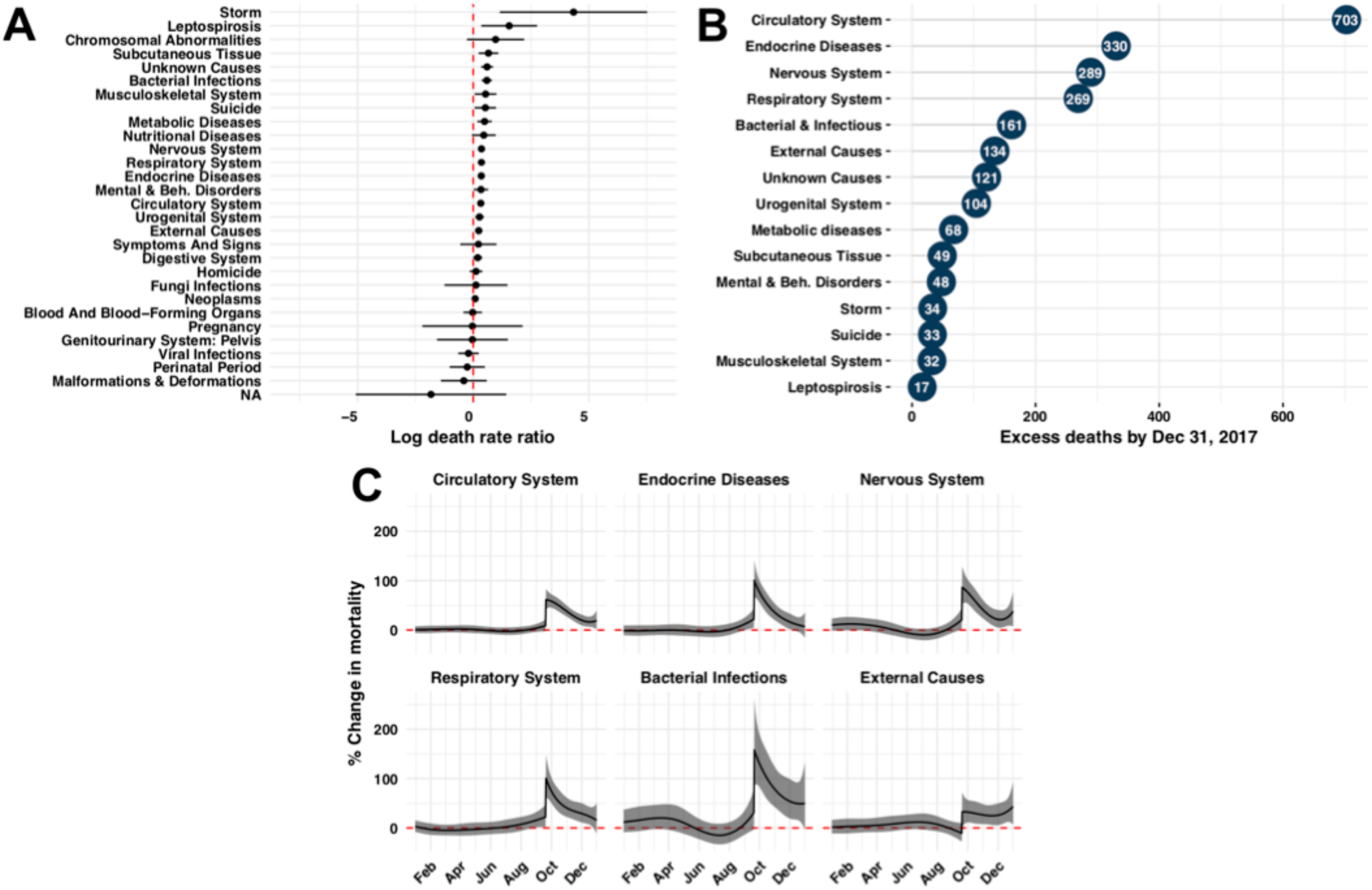
Increase in death rates for 30 causes of death after Hurricane María for the period starting September 20, 2017 and ending on December 31, 2017. A) Log death rate ratio estimates and Bonferroni corrected 95% confidence intervals. B) Estimated excess death by cause of deaths for causes of death with p-values below 0.05 / 30. C) Estimated daily percent increase in death rate after hurricane María for the top six causes of excess deaths: Circulatory System, Endocrine Diseases, Nervous System, Respiratory System, Bacterial Infections, and External Causes. The grey ribbon represents marginal 95% confidence intervals

### Hurricane Sandy (New Jersey)

We obtained monthly death counts for New Jersey from the Underlying Cause of Death database from 2000-2016. We also obtained yearly population estimates from the US Census and interpolated to obtain monthly estimates. (Supplementary Figure 2F)

### Hurricane Harvey (Texas)

We requested daily death counts from Texas’ Vital Statistic System, but our petition was denied. The Underlying Cause of Death database does not have data available for 2017.

### Statistical Methods: Daily counts

We assumed that the death counts *Yi,j* for the *j*-th day of the *i*-th year follow a Poisson distribution with rate:

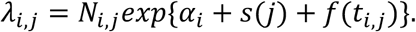

Here *N i,j* is an offset to account for the changing population size, α_*i*_ accounts for the year-to-year variability not due to hurricanes, *s(j)* is the seasonal effect for the *j-*th day of the year, *t*_*i,j*_ = 365 * (*i* - 1) + *j* is time in days, and *f(t*_*i,j*_) accounts for the remaining variability not explained by the Poisson variability. So, for example, a virus epidemic will make *f(t*_*i,j*_) go up slowly, eradication of this epidemic will make *f(t*_*i,j*_) go down slowly, and a catastrophe will make *f(t*_*i,j*_) jump up sharply. We therefore assume *f(t*_*i,j*_) is a smooth function of *t*_*i,j*_ except for the days hurricanes make landfall in which the function may be discontinuous.

Since *s*(*j*) is seasonal, we use Fourier’s theorem and model it as:

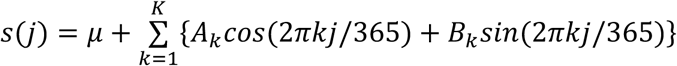

We include an intercept *μ*, which represents the baseline rate for the entire period being studied. We assume that *f(t*_*i,j*_) is a natural cubic spline with *L* equally spaced knots *τ*_*1*_, …, *τ*_*L*_, except that the closest knot to the hurricane day is changed to be exactly on the hurricane day and we permit a discontinuity at this knot. Since natural cubic splines can be represented as a linear combination of basis function and *S(j)* is a linear combination of known functions, ours is a generalized linear model (GLM) and, in theory, we can estimate.α_*i*_, *s*, and *f* using maximum likelihood estimates. However, because we want *f* to be flexible enough to capture relatively high-frequency signals, we instead implement a modular approach that first estimates.α_*i*_ s the *s* and 0, then uses these as offsets to estimate *f* Specifically, we assume that for the non-hurricane years, α_*i*_+ *f(t*_*i,j*_) average out to 0 across years and use this assumption to obtain the MLE for *S(j)*, denoted with 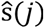. We then use 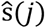. as an offset to estimate the year-to-year deviations α_*i*_ using only months not affected by hurricanes (March-August). We then use the estimate 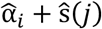 as an offset, estimate *f(t*_*i,j*_) with the MLE and use standard GLM theory to estimate standard errors. Finally, due to lack of data to estimate α_*i*_ for 2018, we extrapolated from the estimates from the previous year (Supplementary Figure 4).

**Figure 4:**
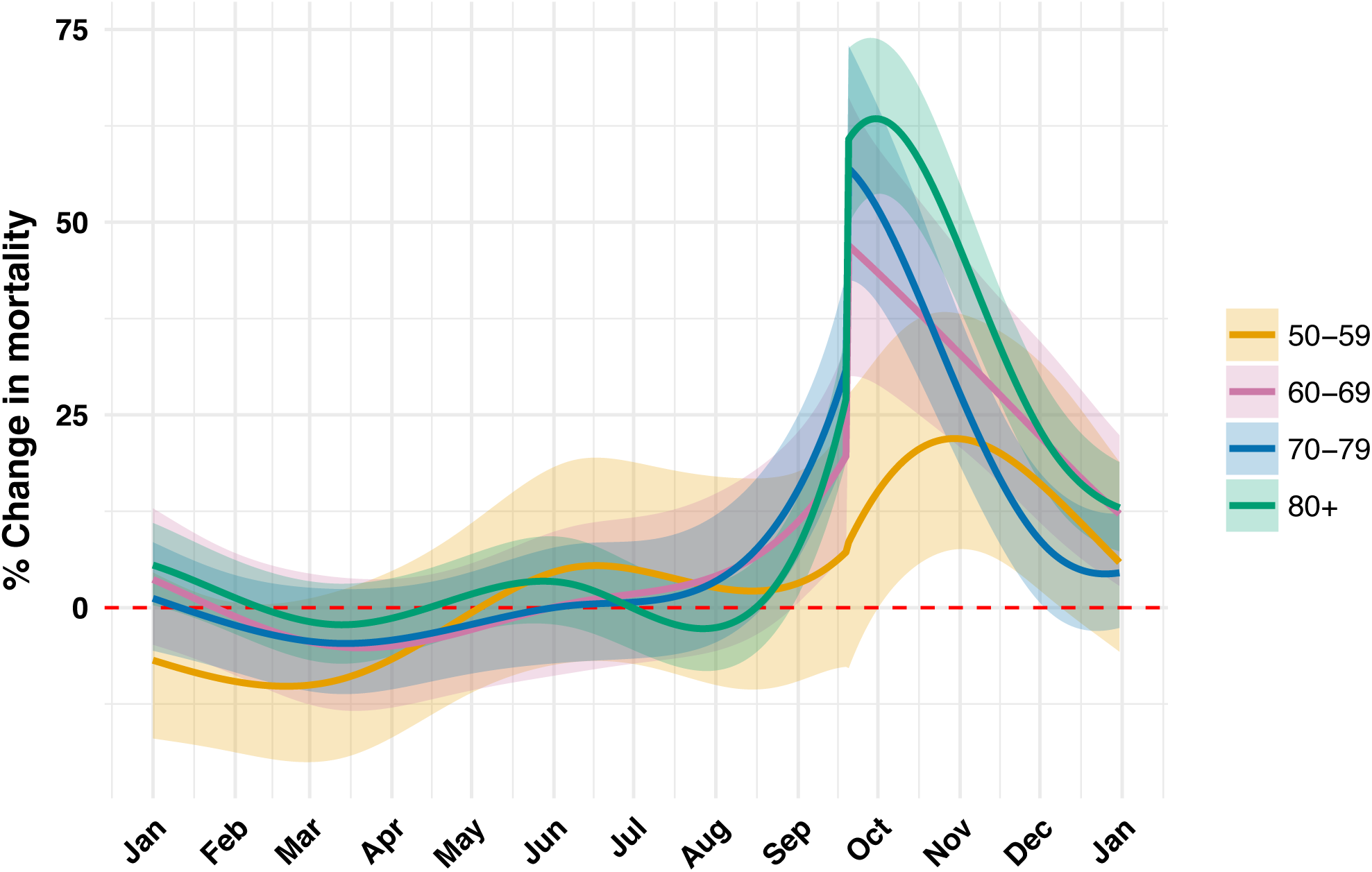
Estimated daily percent increase in death rate by age groups 60-69, 70-79, 80 and higher

For the seasonal effect we use *K=3* since it results in an estimate that captures the general shape of the seasonal trend (Supplementary Figure 5). We used 4 knots per year to model *f(t*_*i,j*_) as this results in a smooth estimate that captures the trend observed in the data (Supplementary Figure 6). We removed years 2001 (Supplementary Figure 6D) and 2014 (Supplementary Figure 6F) for the computation of the seasonal effect: in 2001 there appears to be undercounts in January and in 2014 we see an increased mortality rate in agreement with the Chikungunya epidemic. Diagnostics plots for the residuals, after removing 2011, further show that the model fits the data (Supplementary Figure 7). For the monthly data, we fit a monthly version of this model (See supplementary methods for details).

### Excess death estimates

We first determined the period of indirect effect for each hurricane. To do this, we defined the interval starting on landfall day, denoted here with *t*_0_, until the first day, *t*_*i,j*_> *t*_0_, for which *f(t*_*i,j*_) ≤0. Because we do not observe *f(t*_*i,j*_), and instead obtain an estimate, denoted here with 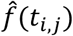, we took the conservative approach of defining the period with the day for which the lower part of a marginal 95% confidence interval crosses 0: 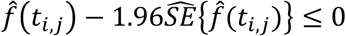. We denoted this interval with *I* and defined the excess deaths by adding the observed deaths minus expected deaths for every day in the interval:

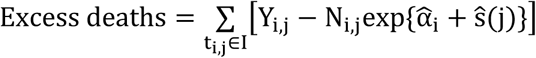

We construct a 95% confidence interval using the following approximation for the Poisson model:

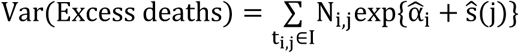

### Natural variability

We quantify the day-specific variability with the observed standard deviation across non-hurricane years for *f(t*_*i,j*_). Because of years like 2001 and 2014, we estimate this standard error with the medial absolute deviation. We refer to this as the *natural variability.*

### Cause of Death

To examine if any cause of death was more prevalent after the hurricane, we used the individual records data spanning 2015-2017. We did not include 2018 data because it appears incomplete (Supplementary Figure 8). We divided causes of death into 30 categories (Supplementary Table 2) and, for each of these, we computed the observed death rate during the September 20 - December 31 period for 2017. We then compared these to the expected rates computed with the 2015-2016 data for that same period. We used the Poisson model to compute confidence intervals. To estimate a daily effect for a cause of death, we fit the Poisson GLM model described above.

### Code

All the code used to implement the procedures described above are included in a GitHub repository: (https://github.com/rafalab/maria). We conducted our analysis using R version 3.4.4.

## Results

### Indirect effects

The death rate in Puerto Rico increased by 73.9% (95% CI, 63.1%-85.4%) the day after Hurricane Maria made landfall (Figure 1, Supplementary Figure 6G) and did not return to the historical average until at least April 15, 2018. During this period, the average increase was 22% (Figure 1, Supplementary Figure 6G). The effects of Katrina were much more direct. On August 29, 2005, the day the levees broke, there were 834 deaths, which translates to an increased in death rate of 689%. However, the increase in mortality rates for the four months following this catastrophe were 17%, 9%, 11%, and 5% percent respectively (Supplementary Figure 9). For Georges, we observed a similar pattern to Maria: a sharp increase to 33% (95% CI, 25%-41%) on landfall day and the death rate not returning to the historical average until January 5, 1999 (Figure 1, Supplementary Figure 6C). The average percent increase in this period was 9%. None of the other hurricanes examined had noticeable indirect effects.

### Excess deaths

We estimated excess deaths of 3,433 (95% CI, 3,189-3,676) for Maria in Puerto Rico, 1,832 (95% CI, 1,600-2,064) for Katrina in Louisiana, and 1,427 (95% CI, 1,247-1,607) for Georges in Puerto Rico (Figure 2). Note that the numbers in the abstract are rounded to the closest 100 for consistency with the precision implied by standard errors of about 100 deaths. These estimates were calculated over periods of 207, 125, and 106 days after landfall respectively. We note that the way in which these deaths accumulated through time were distinctively different. Namely, 39.9% of the excess deaths associated with Katrina occurred on the day the Levee’s broke, while for the Puerto Rico hurricanes, the excess deaths accumulated slowly through a period of months after landfall (Figure 1, Supplementary Figure 6C, Supplementary Figure 6G).

### Cause of Death

Increases in rates were not uniformly seen across all cases of death after Hurricane Maria. Instead, we observed increases for a subset of the causes (Figure 3A). Not surprisingly, storm related deaths showed the largest increase. However, in terms of total excess deaths diseases of the circulatory, nervous, endocrine and respiratory systems explained well over 65% of deaths until at least December 31, 2017 (Table 1, Figure 3B). The indirect effects were for these causes were substantial (Figure 3C).

### Death Rate by Age and Gender

The most affected demographic groups were, from most to least: 80 years and older, 70-79, and 60- 69 and 50-59 with this last group mostly affected by indirect effects (Figure 4). Although younger demographics were not significantly affected (Supplementary Figure S10), these results demonstrate that a large proportion of the population was indeed affected by the indirect effects. We found no significant differences between genders (Supplementary Figure 10).

### Natural variability in excess death rates

The standard deviation of 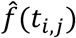 taken across non-hurricane years translated to increases of about 4%, with values as high as 6% in parts of the winter (Supplementary Figure S11A). As a result, for a period of, for example, 103 days (September 20 to December 31) these levels of variability translate into a standard deviation of excess deaths of about 300. For a period of 207 days, the standard deviation grows to about 500 (Supplementary Figure S11B).

## Discussion

On August 29, 2005, a surge due to Hurricane Katrina breached levees and flooded several residential areas in the New Orleans area. This was catastrophic and caused over 600 direct deaths and more to 1,500 indirect deaths in the following four months. In a June 2006 report^16^, the U.S. Army Corps of Engineers admitted that faulty design specifications, incomplete sections, and substandard construction of levee segments, contributed to the damage and $14.5 billion^17,18^ has been invested in constructing stronger levees^18^. On September 21, 1998, Hurricane Georges made landfall in Puerto Rico causing great damage to an already fragile electrical grid. The raw mortality data shows a disturbing increase in mortality rates (Supplementary Figure S12) and using a formal statistical model, we estimated close to 1,400 excess deaths due to this hurricane, an overall impact similar to that of Katrina. In contrast to the response in Louisiana, as far as we know, no systematic effort was put in place to improve Puerto Rico’s electrical grid nor its fragile health system. On the contrary, it appears that the grid continued to deteriorate for the next 19 years^19^. Tragically, after Hurricane Maria made landfall the electrical grid was destroyed leaving 100% of the population^20^, including health facilities, without electricity. It has been well documented that restoration of the electrical grid has been slow^21^, with some estimates reporting that only 30% of the population had electricity a month after the tropical storm^22^. We estimate that, as a result of this fragile health system, a large proportion of the population was affected and as many as 3,000 excess deaths occurred. The insights presented in our analysis should be considered in preparation efforts for the next hurricane.

It is important to note that our estimates and confidence intervals are for the excess deaths above the expected count and that, given the observed variation in non-hurricane years, we can’t attribute all these deaths exclusively to the hurricane, nor can we say that the toll was actually much higher. The same is true for other published estimates^4,9,13^. If we include this natural variability in the uncertainty estimates, the confidence intervals would grow by as much as 1,000 on each limit. This underscores the importance of going beyond analyses based on just monthly counts and historical averages. For example, the sharp jump and then slow decrease in death rate observed on landfall day (Figure 1) provides strong evidence that the observed increases are in fact due to hurricane. Studying specific causes of the excess deaths (Figure 3, Table 1), which do not show increases for causes such as murders, viruses, and cancer, provide further support for the hurricane being the main cause.

In our analysis, the number of knots defining our splines, the number of harmonics in the seasonal effect, and the way we extrapolate to estimate the 2018 death rate were chosen by visual inspection. We therefore performed sensitivity analysis on these choices. Changing the number of knots or the number of harmonics had negligible effects (Supplementary Figure S13 and Supplementary Figure S14). The choice of the expected post Maria death rate did have a larger effect. If we assumed a lower death rate (consistent with unhealthy individuals leaving) the excess death estimate increased and if we assumed a higher death rate (consistent with younger healthier individuals leaving) the excess death estimate decreased (Supplementary Figure S15). We also examined how the estimate changed with different population estimates (Supplementary Figure S15). None of the changes led to a different overall conclusion. The code needed to recreate all the figures and tables is available at: https://github.com/rafalab/maria and we invite others to use our publicly available code and data to try out other approaches.

## Acknowledgments

We thank María M. Juiz Gallego and Jose A. López Rodríguez from the Department of Health of Puerto Rico for diligently providing all the data we requested. We thank the Puerto Rico Institute of Statistics for providing population data. We thank Canay Deniz, Andrea Samdahl, Lara Montini and Ilya Vasilenko from Teralytics for sharing their data and providing helpful explanations. We thank Matthew Kiang for many valuable demography insights and Deepak Lamba Nieves for suggesting readings on Puerto Rico’s electrical grid. We thank Raul Figueroa for critiques that led us to perform the sensitivity analysis. Finally, we thank all the authors of NEJM paper whose publication appears to have resulted in the government releasing the data. In particular, we thank Caroline Buckee for getting it all started.

This findings and conclusion are those of the authors and do not necessarily represent the official position of the Florida Department of Health.

## Supplementary Materials

### Supplementary Methods

#### Statistical Methods: Monthly counts

For the monthly data, we fit a monthly version of the model above. Because the counts are much larger once we aggregate at this level, we made use of the normal approximation to the Poisson. Specifically, we defined the monthly rates as 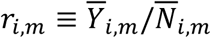 where 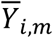 is the average number of deaths in month *m* of year *i* and 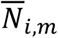 is the person years for that period. Thus, we collapsed the model above to a monthly version as follows: we assumed that the monthly rates can be described with the following model:

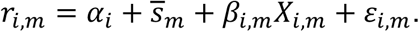

Here α_*i*_ accounts for year-to-year variability as in daily data model and 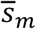 is the average of.*s(j)* for all days *j* in month *m*. Because we no longer need splines to model the effects, we instead use indicator functions to denote if a month/year was affected by the hurricane. Specifically, we define *X*_*i,m*_ as an indicator that is 1 for the months) in year *m* that were affected by the hurricane and 0 otherwise. The parameter *β*_*i,m*_ thus represents the effect of the hurricane on death rate and is equivalent to the integral of *f(t*_*i,j*_) for *t*_*i,j*_ in month *m* of year *i* The natural, yet non-hurricane related variability, is represented by the term *ε*_*i,m*_ which are assumed to be independent and normally distributed with average 0 and month-specific standard deviation *σ*_*m*_. Notice that this is a standard linear model and the estimates can be obtained with least squares.

**Figure S1:**
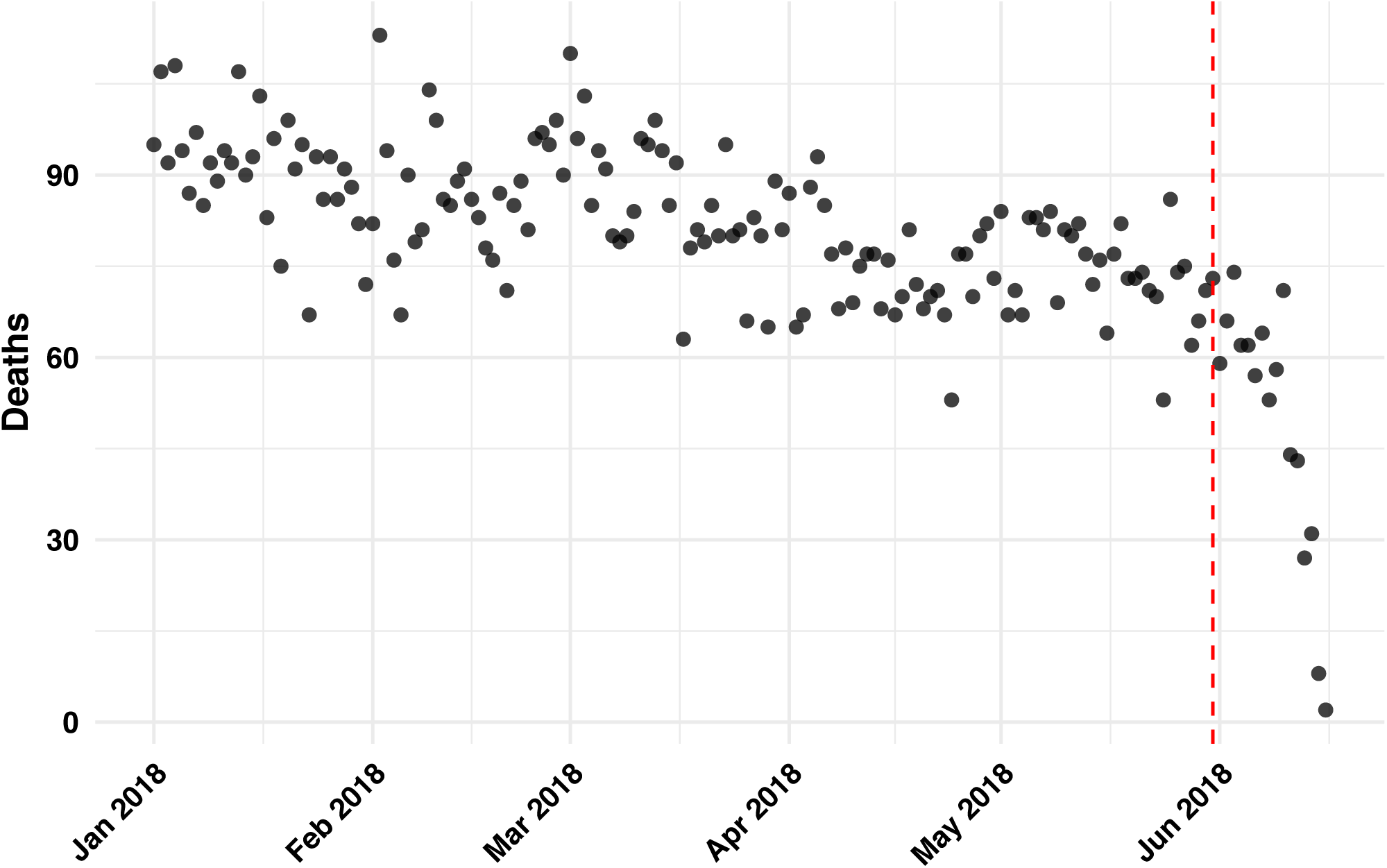
Reported deaths by day for Puerto Rico daily data in 2018. We excluded data to the right of the vertical dashed line

**Figure S2:**
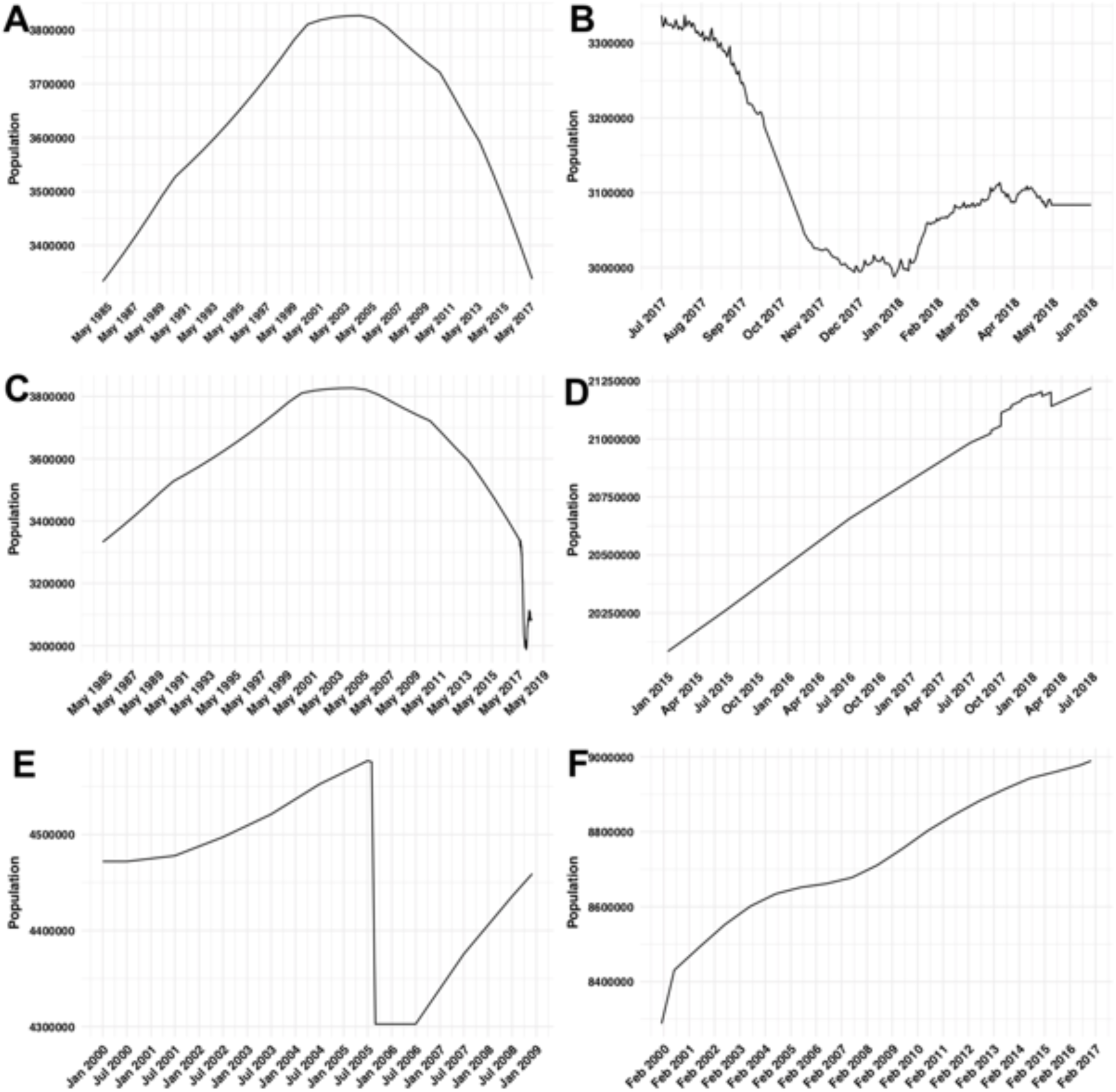
Estimated population sizes. A) Interpolation of yearly estimates for Puerto Rico from 1985 to 2017. B) Estimates for Puerto Rico population from July 1, 2017 to June 1, 2018 based on proportion of population decreased provided by Teralytics Inc. C) Final estimate for Puerto Rico population. D) Estimated population for Florida. Includes increases based on Teralytics Inc. estimate of immigration from Puerto Rico. E) Estimated population for Louisiana based on interpolation and extrapolation. F) Estimate population for New Jersey based on interpolation

**Figure S3:**
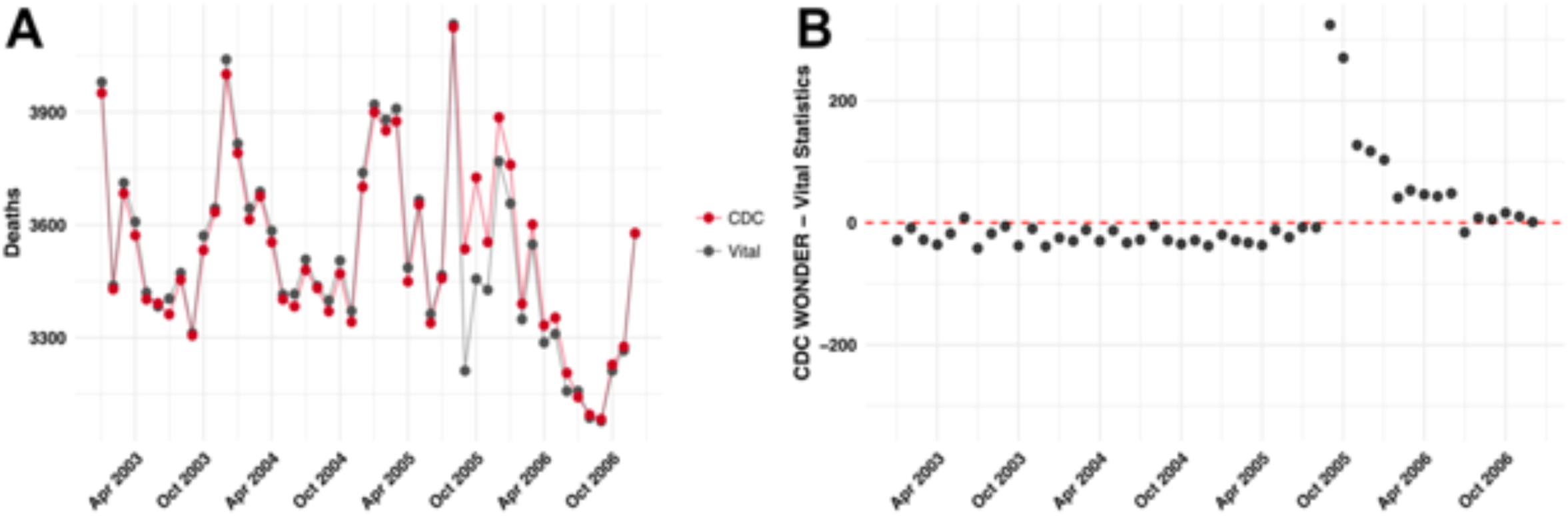
Comparison of census monthly death counts to Vital Statistics System monthly death counts. A) Total death counts. B) Difference between estimates

**Figure S4:**
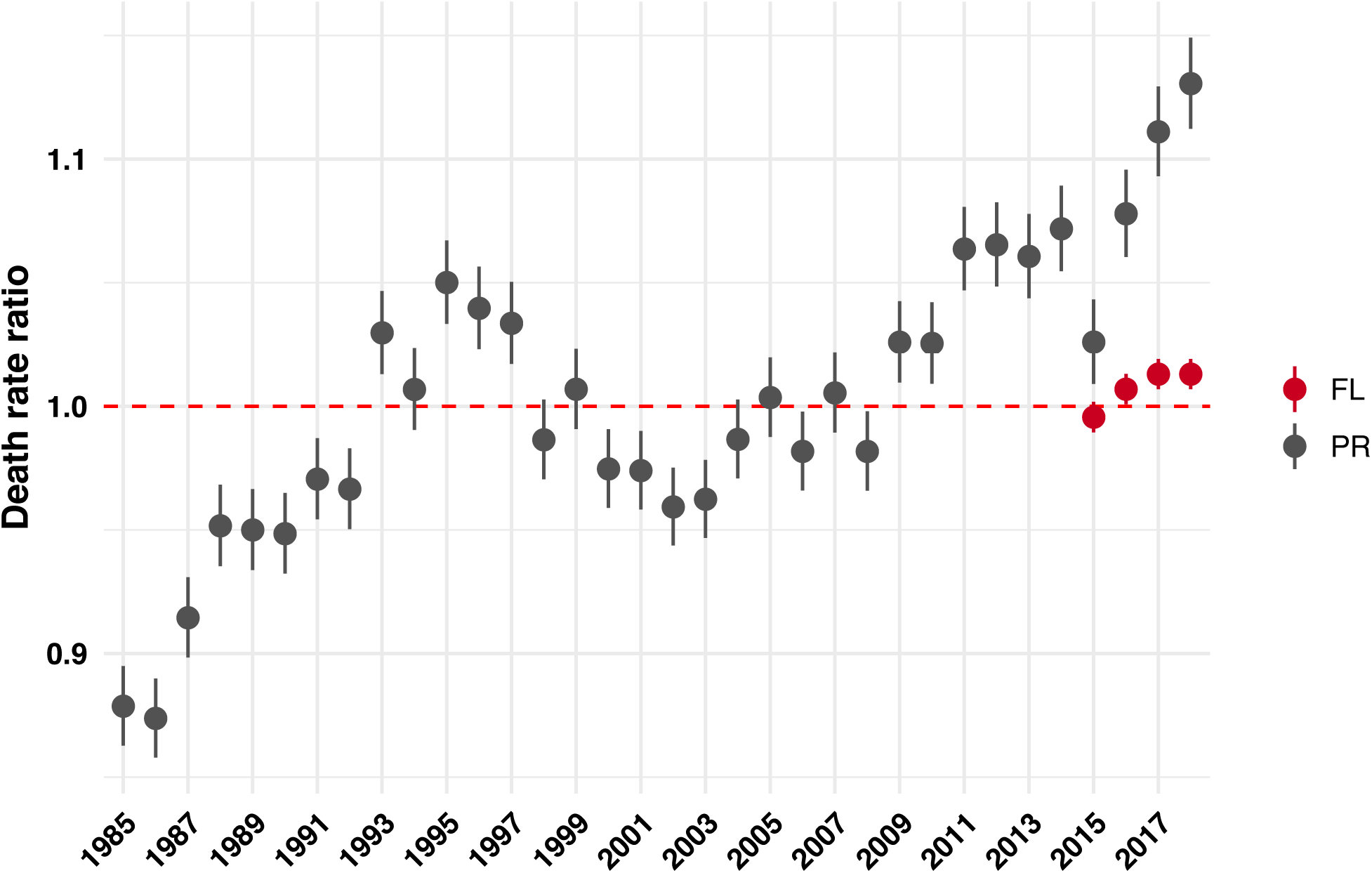
Estimated yearly offsets and marginal 95% confidence intervals for Puerto Rico and Florida obtained from fitting the GLM. The 2018 data is based on extrapolation obtained by fitting a natural cubic spline to the rest of the data for Puerto Rico and by assuming the same value as 2017 for Florida

**Figure S5:**
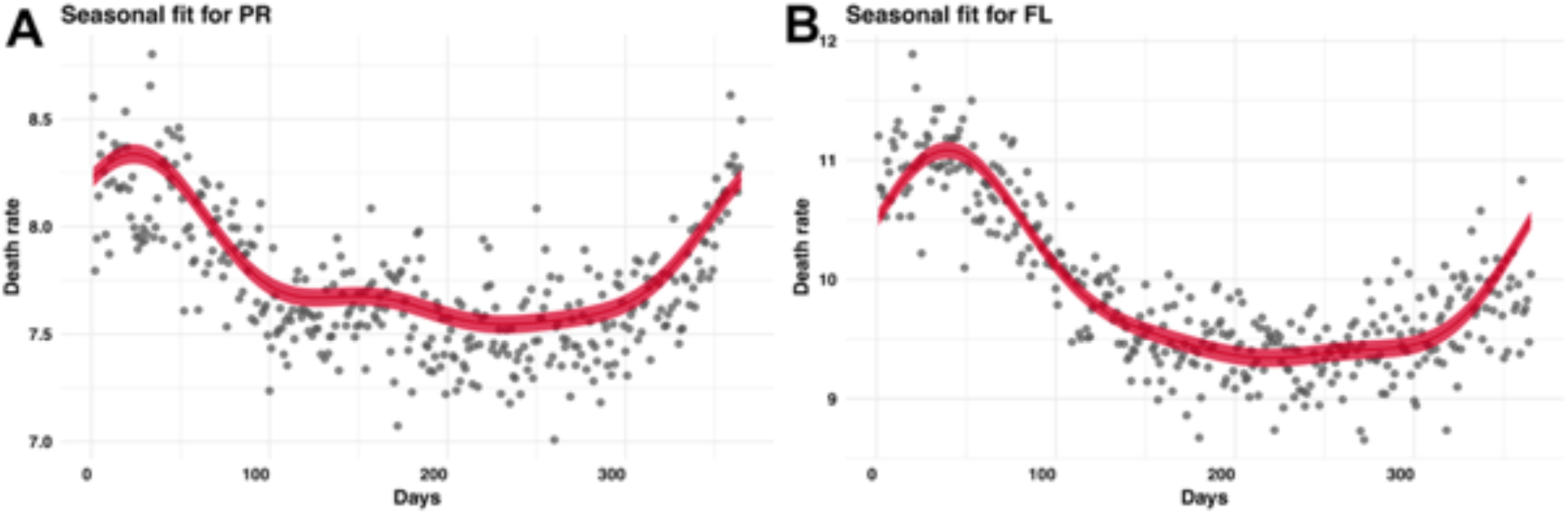
Estimated seasonal effects. Estimated seasonal effects from the GLM and marginal 95% confidence intervals. A) Estimates for Puerto Rico. B) Estimates for Florida

**Figure S6:**
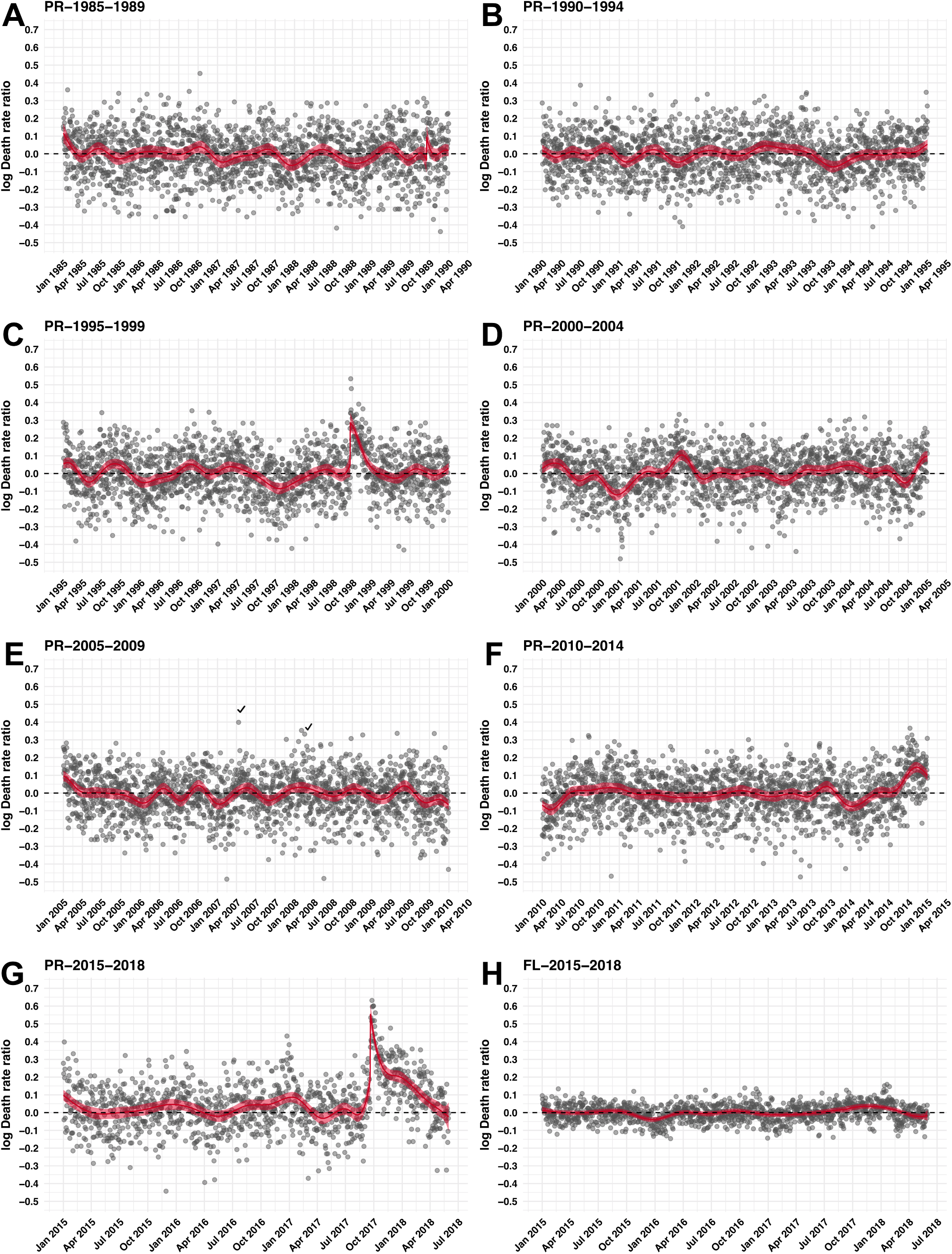
Model fit check. We compute the residuals after fitting the seasonal and yearly effects and plot these against time. The fitted *f(t_i,j_)* and marginal 95% confidence interval are shown to capture trends. A) Puerto Rico 1985-1990 (includes Hugo). B) Puerto Rico 1990-1995. C) Puerto Rico 1995-2000 (includes Georges). D) Puerto Rico 2000-2005. E) Puerto Rico 2005- 2010. F) Puerto Rico 2010-2015 (Includes Chikungunya epidemic). G) Puerto Rico 2015-2018 (includes María). H) Florida 2015-2018 (includes Irma)

**Figure S7:**
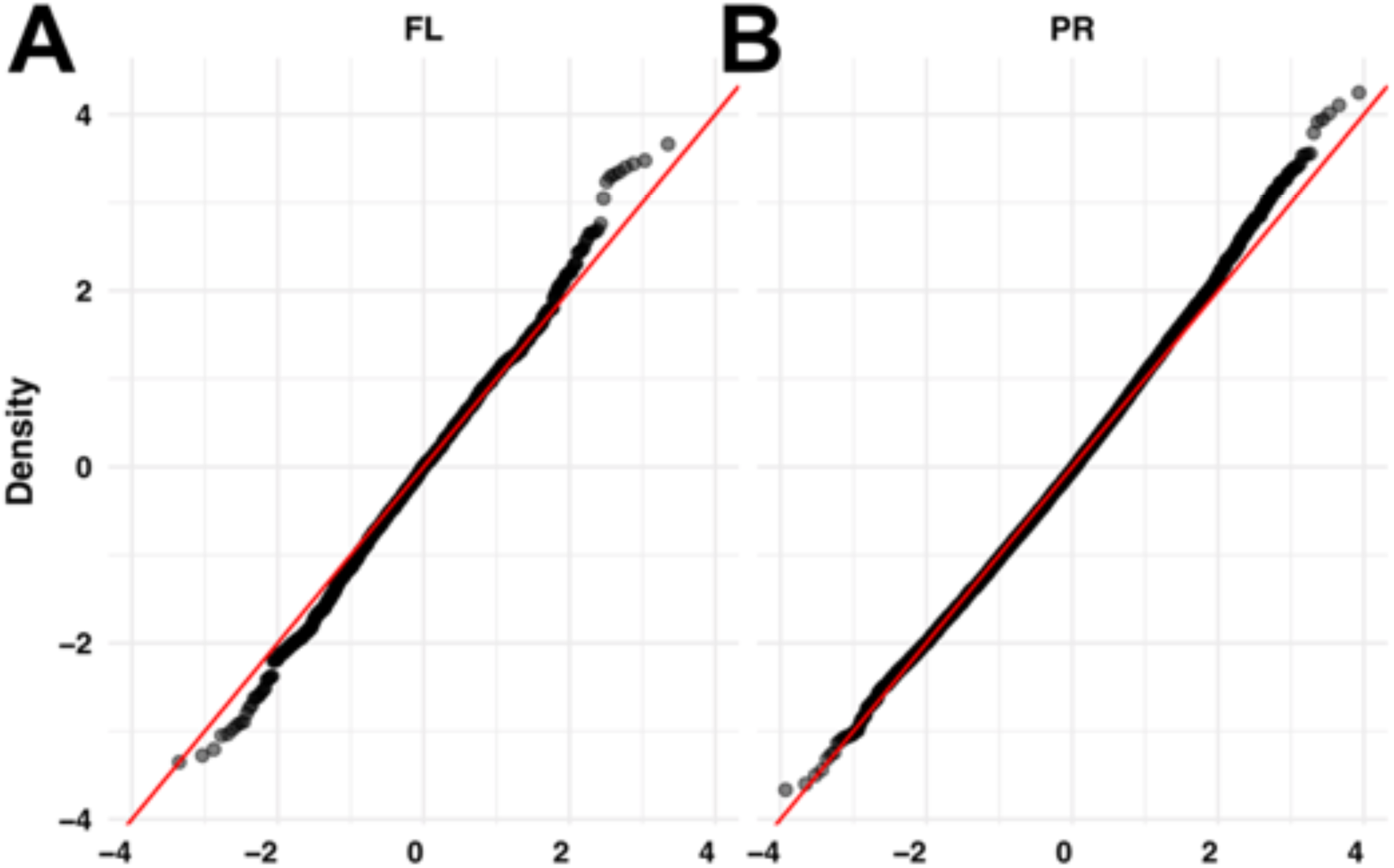
Poisson assumption check. We compute Pearson residuals and generate a normal qq-plot. We remove years determined to be outlier years. A) Florida. B) Puerto Rico

**Figure S8:**
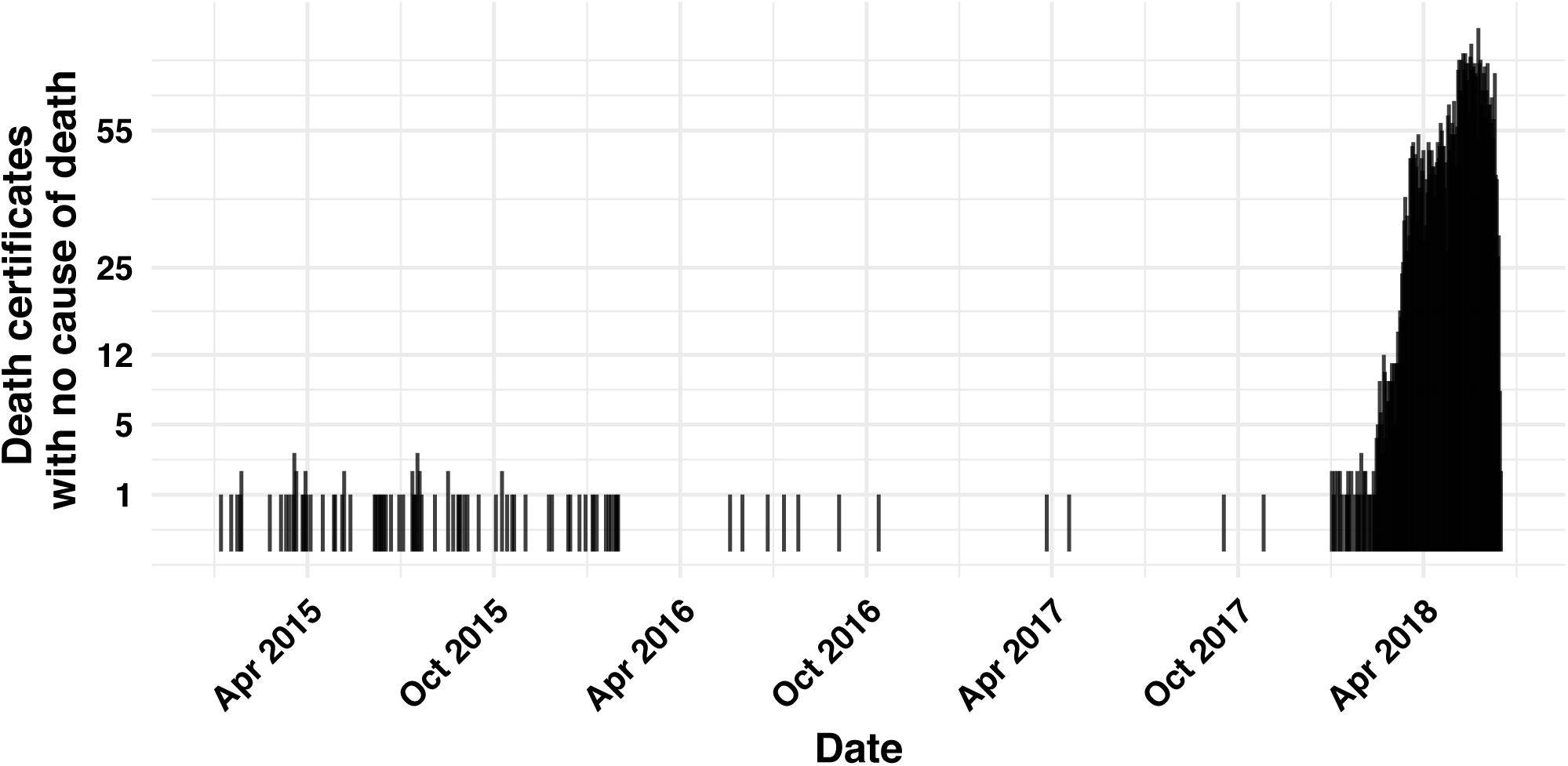
Number of individual records with no cause of death entered by day for the Puerto Rico individual 2015-2018 individual date

**Figure S9:**
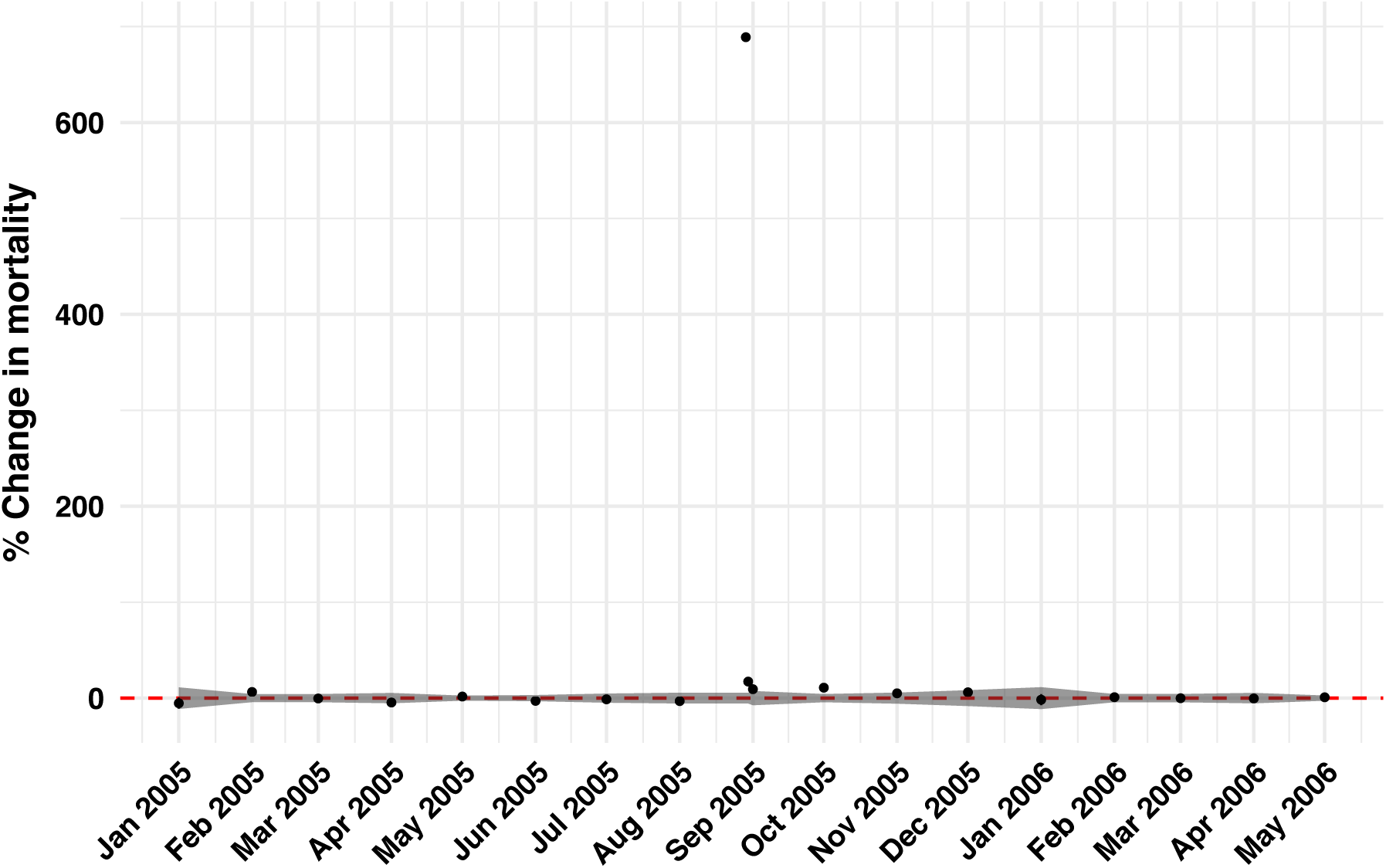
Estimated percent increases in death rates after hurricanes Katrina. Monthly estimates are provided except for August 29, 2005 and August 30-31, 2005 for which daily data was used to compute the number of deaths. The grey ribbon represents the range of variation (plus or minus two standard deviations) in death rate increases seen across non-hurricane

**Figure S10:**
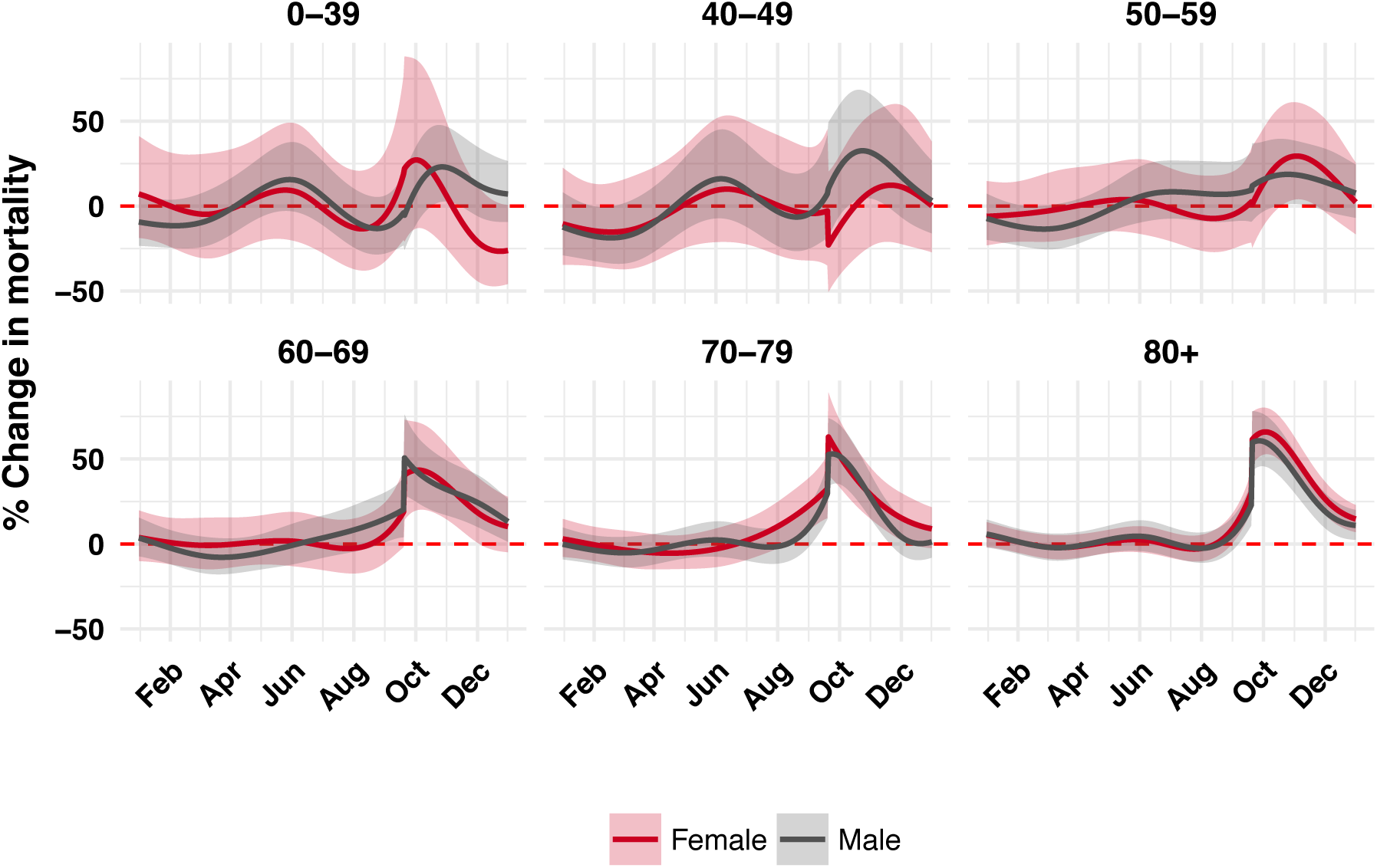
Percent change in mortality by six age groups and by gender.

**Figure S11:**
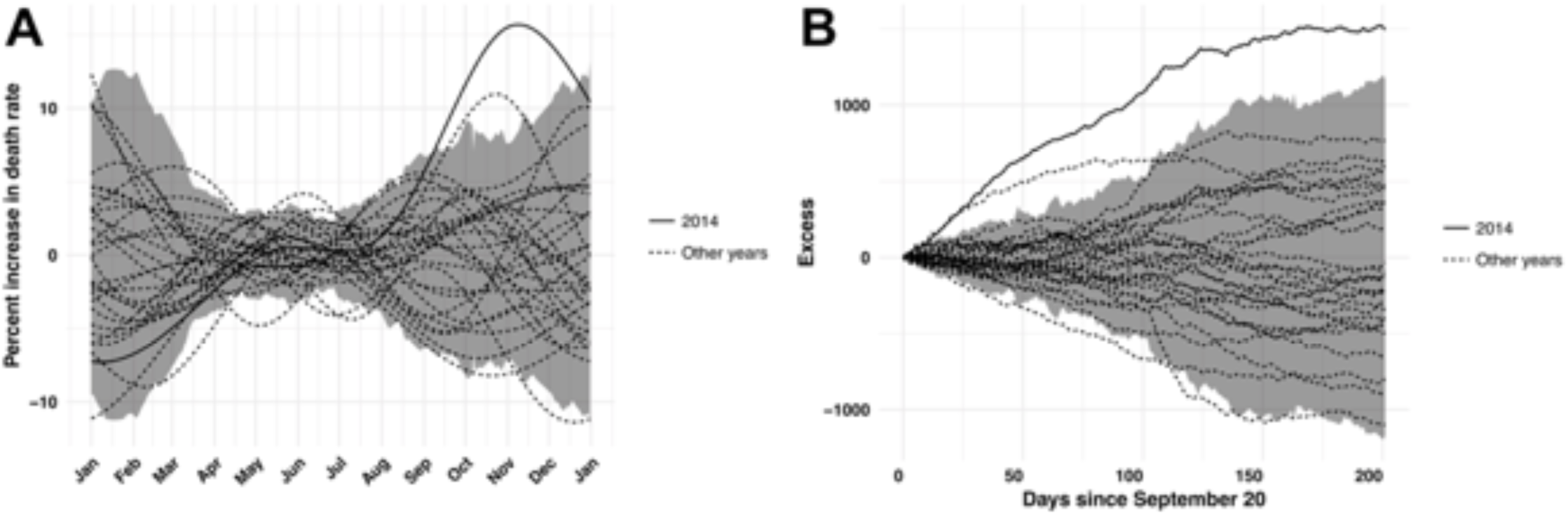
Natural non-hurricane related variation. For each day we estimated the standard deviation using median absolute deviation of *f(t*_*i,j*_) The dashed lines are the *f(t*_*i,j*_) and the grey ribbon is plus or minus two standard deviation

**Figure S12:**
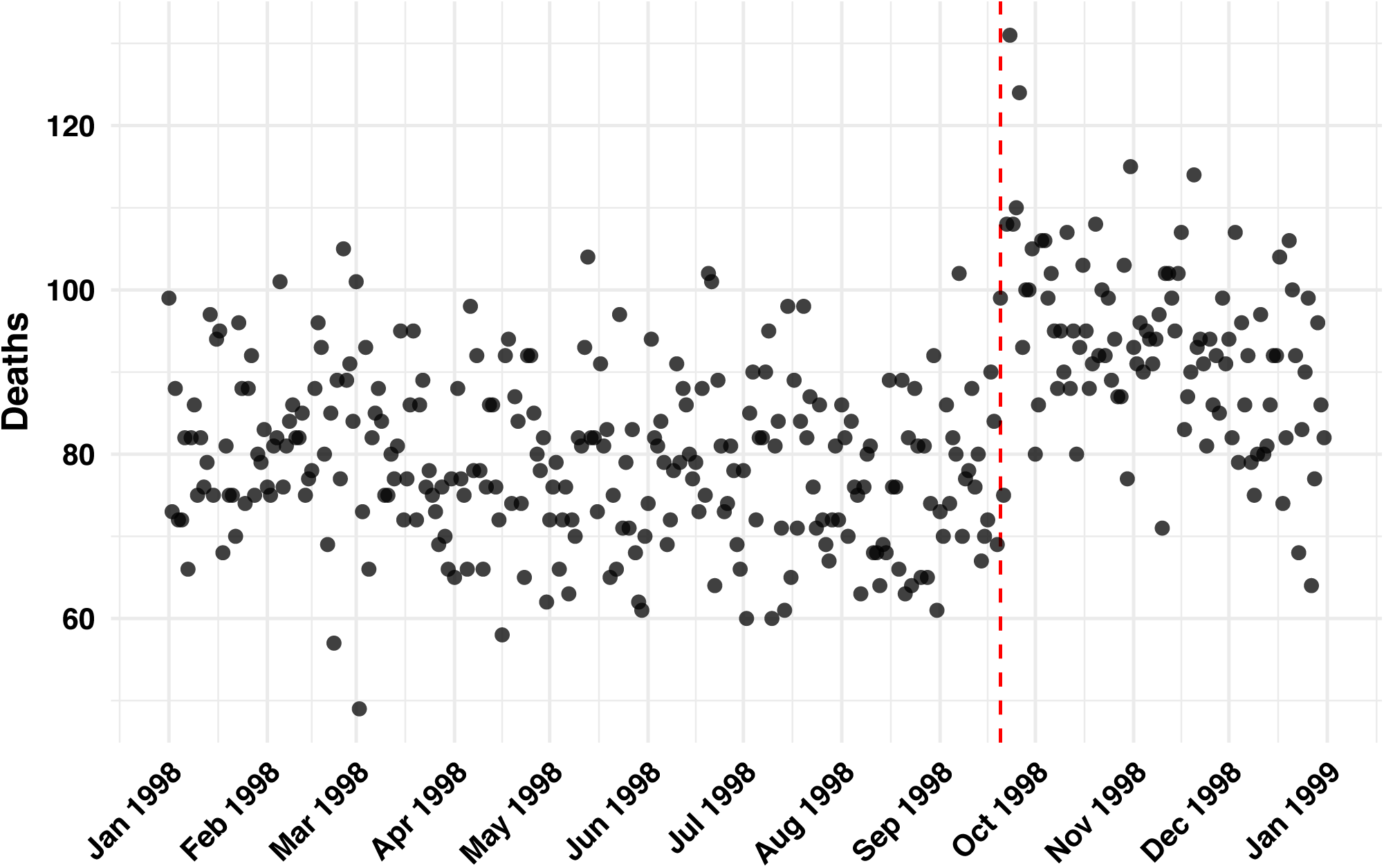
Raw Puerto Rico death count data from 1998, the year Georges hit the island

**Figure S13:**
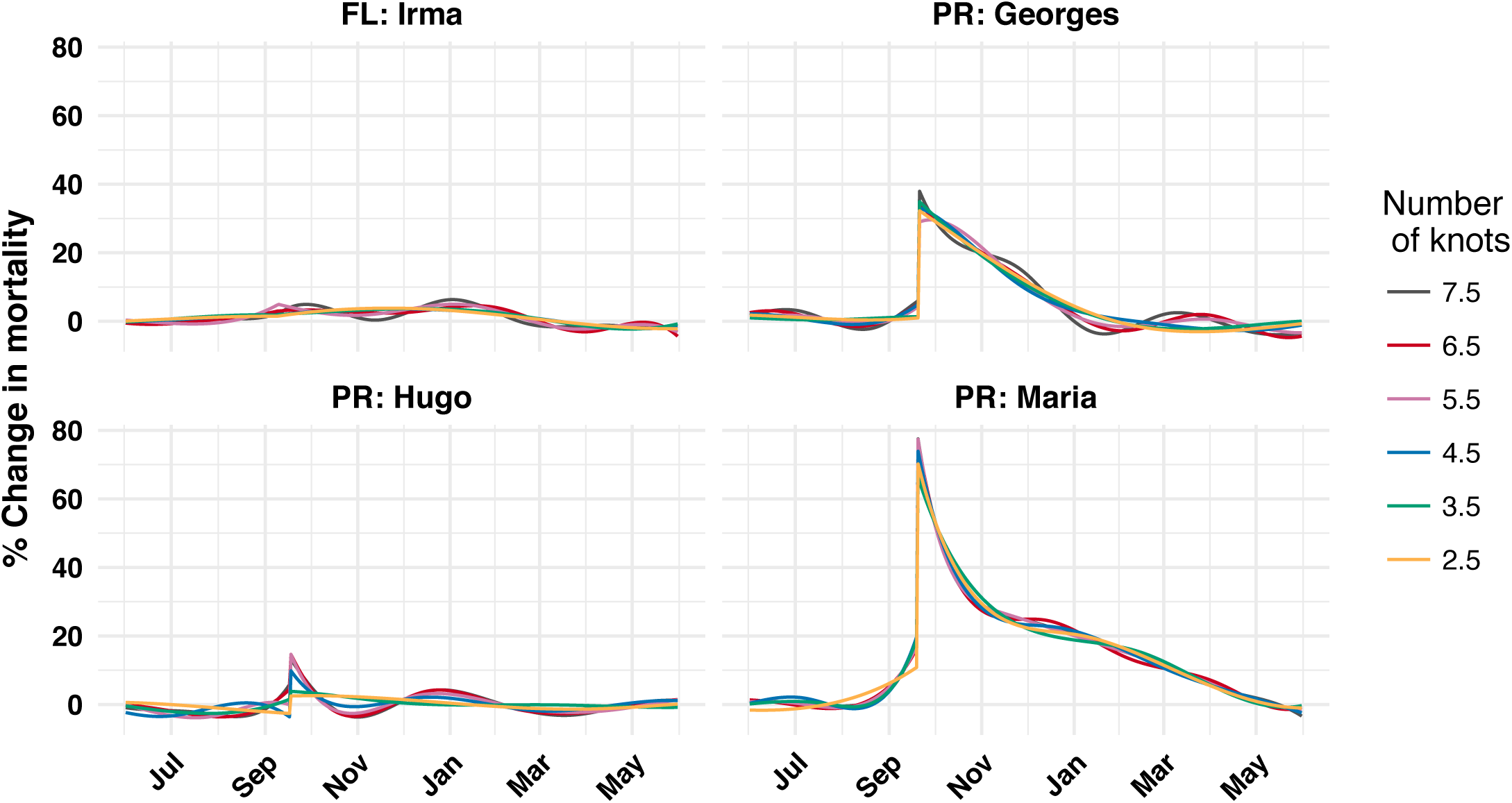
Model fit with different number of knots

**Figure S14:**
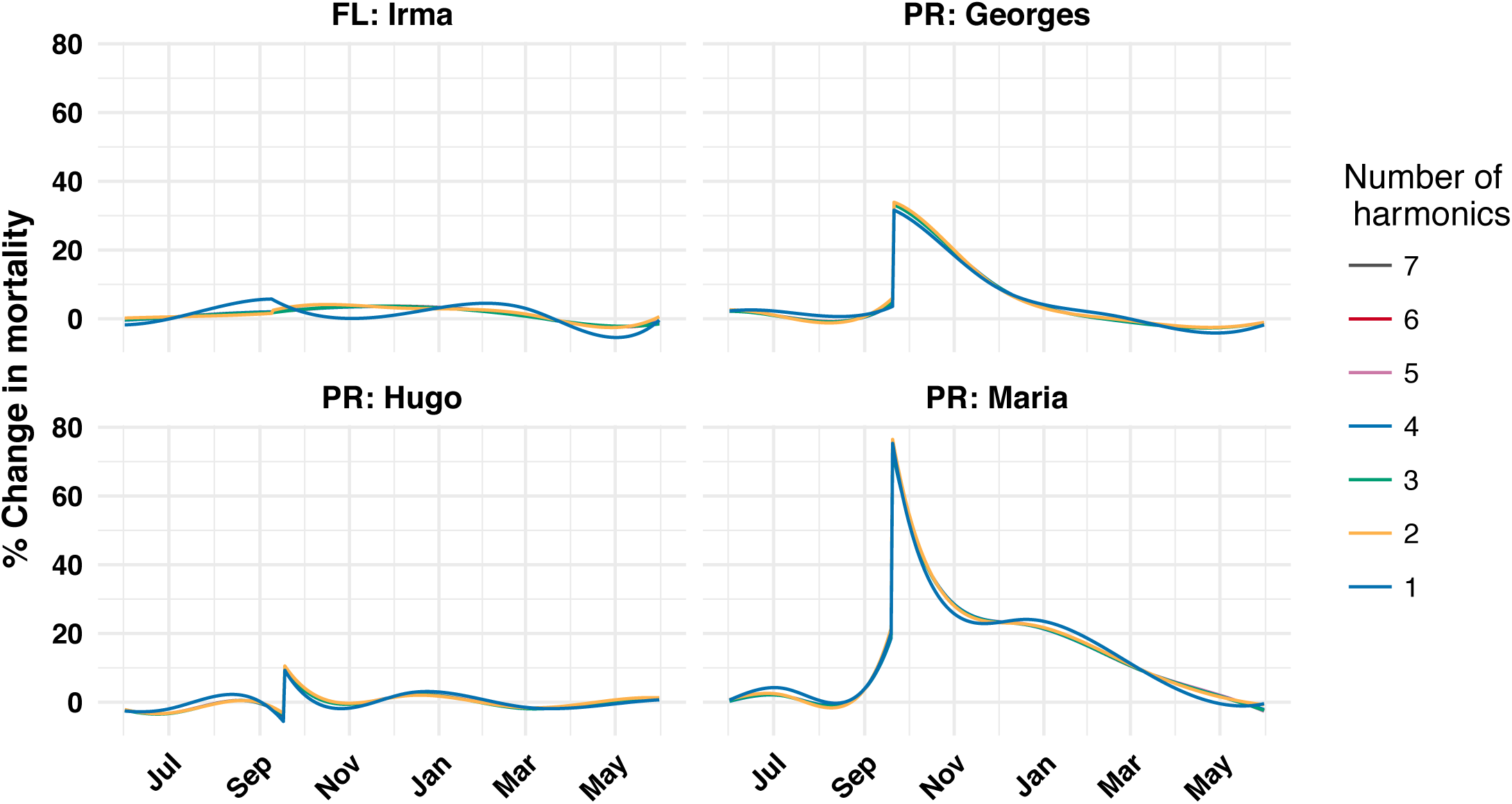
Model fit with different number of harmonics

**Figure S15:**
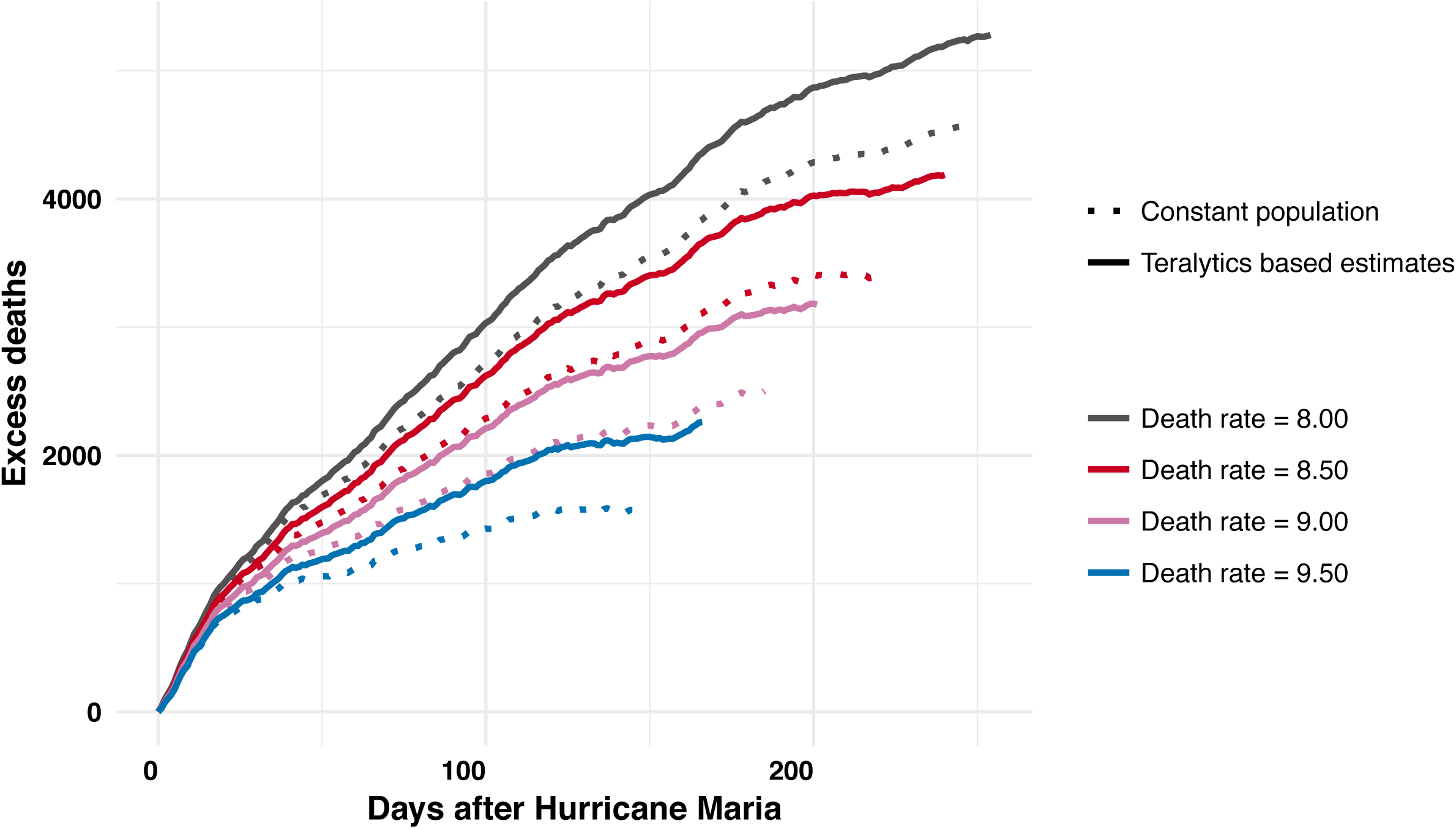
Assuming different post hurricane expected death rates & population estimates

**Table S1:**
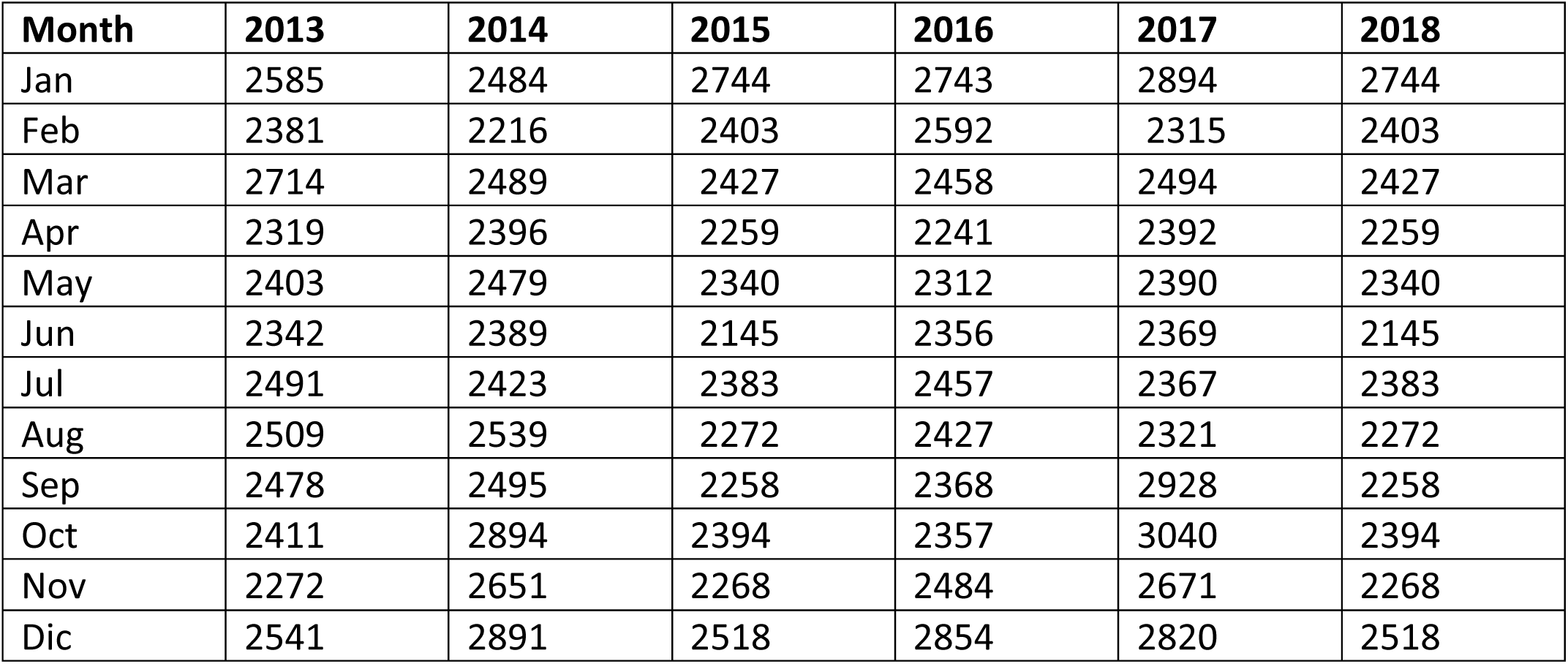
Monthly data released by the government of Puerto Rico after the publication of the *Harvard Study*

**Table S2:**
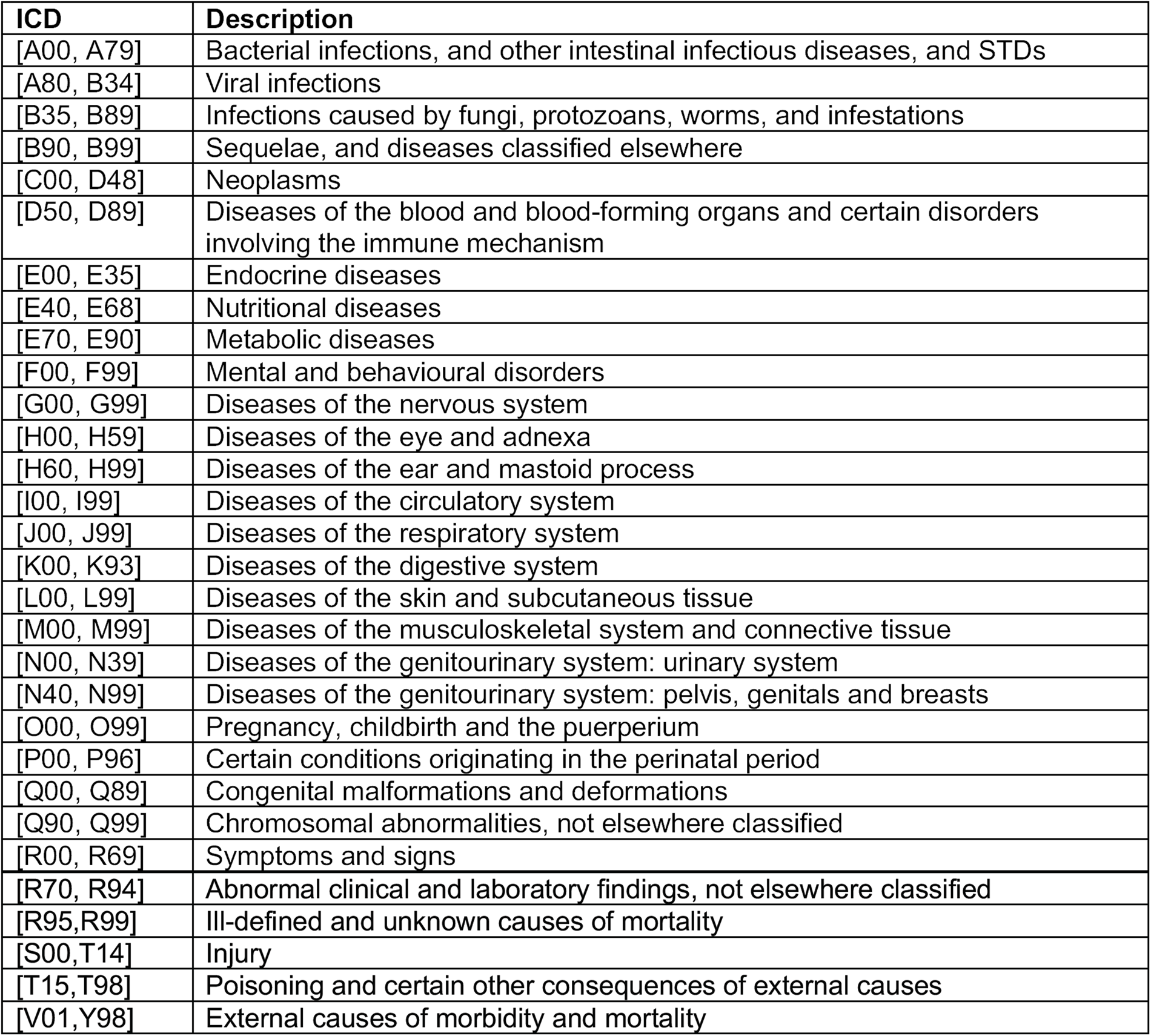
Grouping of causes of deaths

